# The expression tractability of a biological trait

**DOI:** 10.1101/278770

**Authors:** Li Liu, Jianguo Wang, Jianrong Yang, Xionglei He

**Author notes:** Equal contribution (Li Liu, Jianguo Wang). Correspondence to: Xionglei He, School of Life Sciences, Sun Yat-sen University, Guangzhou 510275, China.

## Abstract

Understanding how gene expression is translated to phenotype is central to modern molecular biology, but the success is contingent on the intrinsic tractability of the specific traits under examination. However, an *a priori* estimate of trait tractability from the perspective of gene expression is unavailable. Motivated by the concept of entropy in a thermodynamic system, we here propose such an estimate (*S*_T_) by gauging the number (*N*) of different expression states that underlie the same trait abnormality, with large *S*_T_ corresponding to large *N*. By analyzing over 200 yeast morphological traits we show that *S*_T_ is constrained by natural selection, which builds co-regulated gene modules to minimize the total number of possible expression states. We further show that *S*_T_ is a good measure of the titer of recurrent patterns of an expression-trait relationship, predicting the extent to which the trait could be deterministically understood with gene expression data.

## Introduction

A complex system is characterized by the microscopic configuration of its constituents (or microstate) and the macroscopic property of the system (or macrostate) (Ladyman et al. 2013). Cell is a typical complex system in which the expression of genes represents microstate and cellular phenotype represents macrostate (Komili and Silver 2008). Disentangling the relationships between expression microstate and phenotypic macrostate has been the aim of numerous studies in molecular cell biology, with a focus recently on the production of large-scale data and the development of sophisticated analyzing methods (Janes et al. 2005; Segal et al. 2005; Lee et al. 2008; Zhu et al. 2008; Ayroles et al. 2009; Chen et al. 2009; Civelek and Lusis 2014; Gamazon et al. 2015; Ritchie et al. 2015; Gusev et al. 2016). Despite enormous achievements, the availability of many methods *per se* indicates the limitation of the data and the lack of generality of the methods (Amin et al. 2014; Lloyd et al. 2015; Volm and Efferth 2015), highlighting the fact that the success of the extrinsic research efforts is contingent on the intrinsic tractability of the focal relationships. The importance of knowing *a priori* the tractability of a question to be addressed is widely recognized in mathematics, physics, and chemistry (Ayoub 1982; Feldman and Crutchfield 1998; Allu and Oprea 2005; Lopezruiz et al. 2010). For example, the competition for finding a general formula for solving equations of degree five had lasted ∼300 years since the successes in equations of degree three and degree four in the 16^th^ century, and ended in the 19^th^ century when the impossibility of such a general formula was proved; the subsequently developed group theory that gives an effective criterion for the solvability of all polynomial equations formed the basis of modern algebra (Ayoub 1982; Rosen 2016). However, biology as a branch of science dominated by empirical solutions has seldom been guided by such wisdoms.

## Results

We reason that here a critical issue is, for a given phenotypic macrostate, how many expression microstates there could be. Similar to the situation in thermodynamic systems, a large number (*N*) of possible microstates would suggest a strong uncertainty underlying the given macrostate, understanding of which may rely on stochastic thinking more than the prevailing deterministic thinking in biology. In contrast, straightforward understandings would be readily available if the *N* is small. In research practice each time often a particular trait, which represents a single dimension of the multidimensional macrostate of the system, is examined. For a trait considered qualitatively (e.g., normal versus abnormal), the information flow from genotypes to the same phenotype could be mediated by either the same or different expression microstates (Fig. 1A). Here, the expression state of an individual gene can be categorized as 0, 1, and −1 for no expression difference, up-regulation, and down-regulation relative to a reference level, respectively. If the trait abnormality is described quantitatively, individuals with similar trait values can be treated as qualitatively the same.

**Figure 1.**
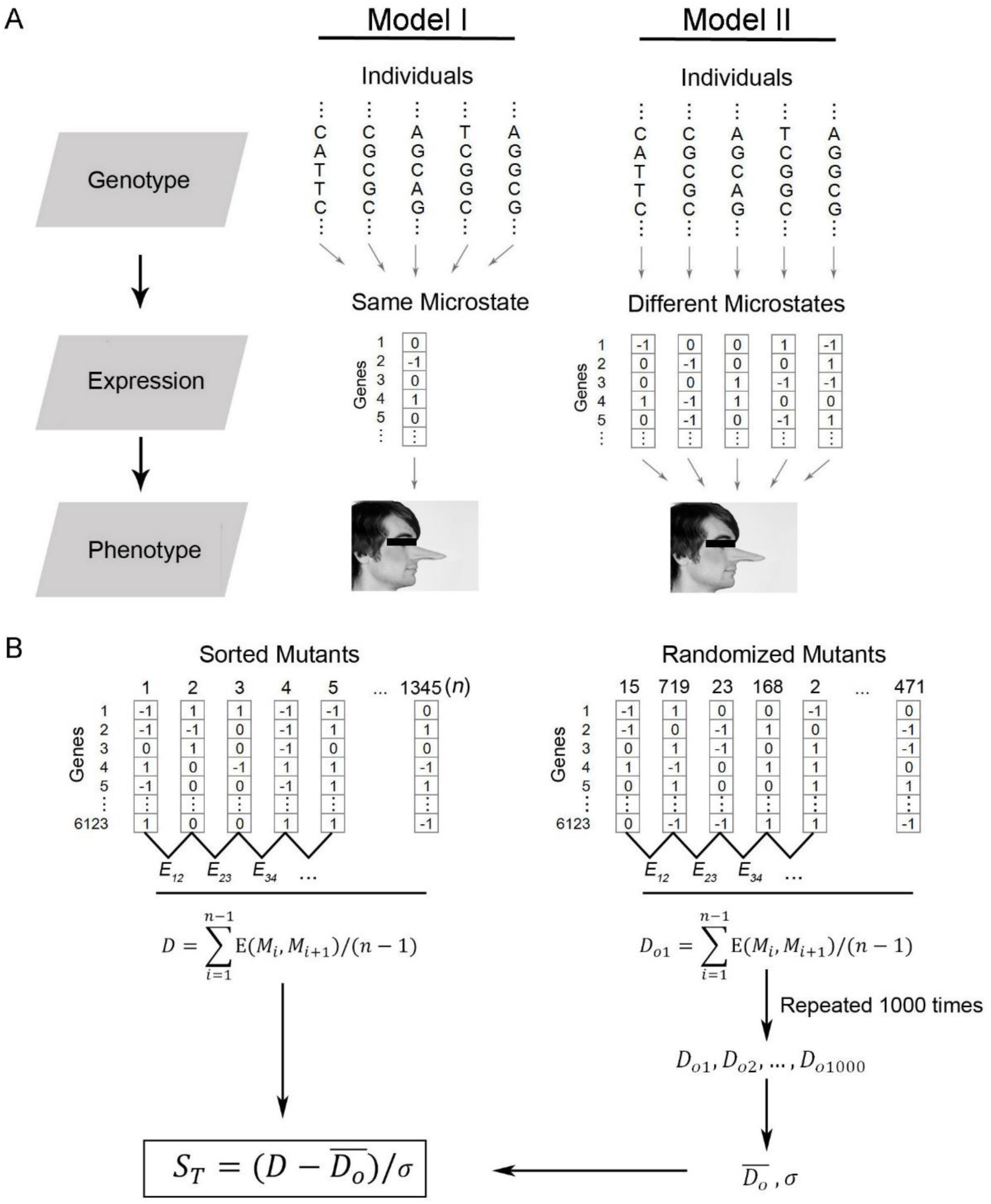
**(A)** Two models of the information flow from genotype to phenotype that differ in the number of the intermediate expression microstates. **(B)** The workflow of calculating *S*_T_ using the 1,345 yeast mutants.

## Measure the divergence of the expression programs underlying a trait

We propose a measure (*S*_T_) of the number of different expression states that underlie the same trait abnormality. Previous studies measured quantitatively 216 morphological traits (e.g., cell size, bud growth direction, area of nucleus region, and so on) of 1,345 yeast *Saccharomyces cerevisiae* gene-deletion mutants whose genome-wide expression profiles are also available (Supplemental Table 1) (Materials and Methods), permitting us to study the expression microstates of a large number of traits simultaneously (Ohya et al. 2005; Kemmeren et al. 2014). For each morphological trait we ranked the 1,345 yeast mutants in an ascending order according to the focal trait values (Fig. 1B). Because of the large number of mutants assessed, neighbouring mutants often showed highly similar trait values (Supplemental Fig. 1). We then computed the average expression divergence (*D*) between two neighbouring mutants using the formula:

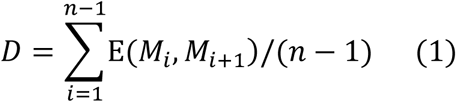

, where *n* (=1,345) is the total number of mutants, *M*_*i*_ and *M*_*i+1*_ represent the mutants ranked *i* and *i+1*, respectively, and *E* is the Euclidean distance of the expression profiles of the two mutants (Materials and Methods). Note that here neighbouring mutants had commonality only in one dimension (i.e., the focal trait) of the phenotypic macrostate. To gauge the expression divergences that represent dimensions independent from the focal trait, we computed *D*_*o*_, which is the *D* of the *n* mutants that are randomly sorted. By repeating the randomization of the mutants 1,000 times we obtained the mean 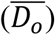 and standard deviation (σ) of D_o_, and then had:

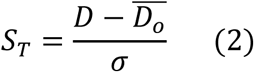

Thus, by controlling for the expression divergence unrelated to the focal trait *S*_T_ measured specifically the divergence of the expression programs underlying the (nearly) same trait alterations.

Using this formula we obtained *S*_T_ for each of the 216 yeast traits (Supplemental Table 2). It varied substantially among traits, ranging from ∼ −7 to ∼2 (Fig. 2A). Because *S*_T_ was effectively the Z-score transformation of *D* in the distribution of *D*_*o*_, −2< *S*_T_ < 2 meant no significant difference between *D* and *D*_*o*_ at the level of *P* ≈ 0.05. Thus, −2 < *S*_T_ < 2 suggested no higher than expected expression convergence underlying the same/similar trait alterations; in contrast, *S*_T_ < −2 suggested that the same/similar trait alterations of different mutants be mediated by shared expression programs. Notably, the observation −2 < *S*_T_ < 2 cannot be explained by strong trait dissimilarity between neighbouring mutants (Supplemental Fig. 2). The across-trait variation of *S*_T_ was insensitive to the number and the identity of the mutants examined, suggesting that *S*_T_ represent an intrinsic feature of the traits (Fig. 2B).

**Figure 2.**
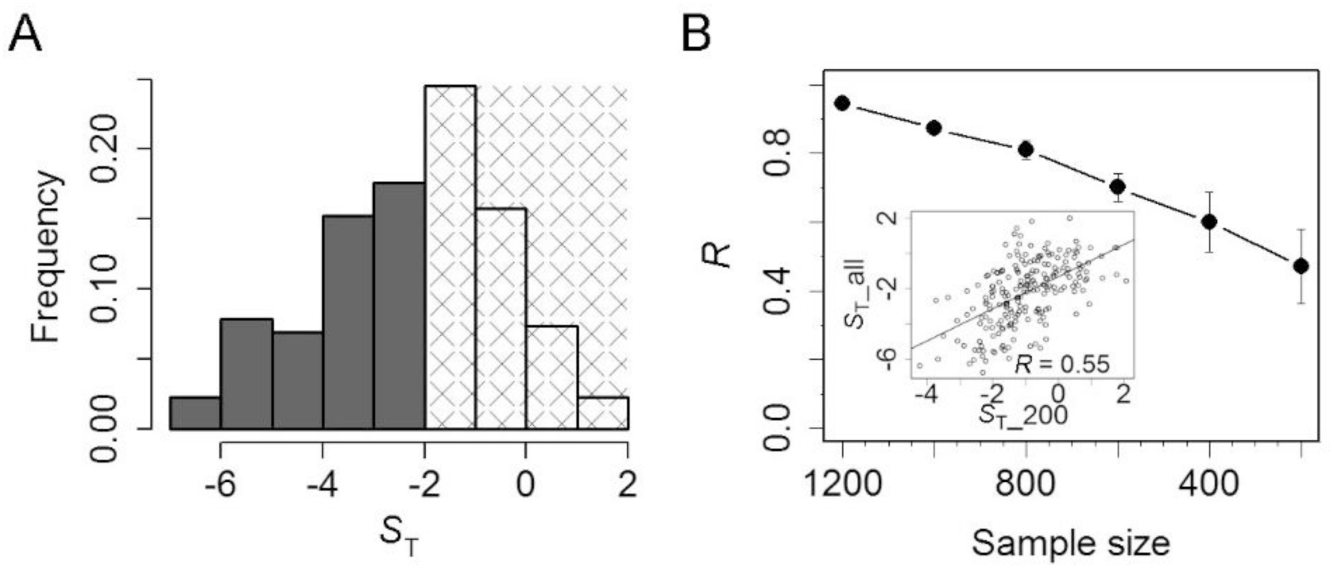
**(A)** Distribution of *S*_T_ of the 216 yeast morphologic traits. Approximately 40% of the traits show −2 < *S*_T_ < 2, a range indicating no higher expression similarity than expected by chance between mutants of similar trait values. **(B)** The relative *S*_T_ of the 216 traits is insensitive to the reduced number of mutants included. Pearson’s *R* between *S*_T_-all based on all 1,345 mutants and *S*_T_N_ based on randomly sampled *N* mutants, where *N* = 1200, 1000, 800, 600, 400, and 200, respectively, is shown. Ten rounds of random sampling are considered for each *N*, and the mean Pearson’s *R* is plotted with the error bar showing one standard deviation. The comparison of *S*_T_-all and *S*_T_-200 derived from one random sampling is shown as an inset.

## *S*_T_ is shaped by natural selection that builds co-regulated gene modules

It is interesting to know what determines the *S*_T_ of a trait. We noted that *S*_T_ is analogous to the entropy of a thermodynamic system. The second law of thermodynamics predicts that the entropy of a thermodynamic system always increases over time unless external forces are applied to the system. We reasoned that natural selection is the only possible external force able to constrain a biological system. We simulated how selection could shape an expression network that underlies traits by considering a previously proposed gene-trait architecture (Chen et al. 2016) (Materials and Methods). The results suggested two insights: 1) Co-regulated gene modules can dramatically reduce *S*_T_; and 2) such co-regulations tend to evolve among genes responsible for evolutionarily more important traits (Supplemental Fig. 3). This is because that the number of possible expression states could be astronomical if the involved genes lack co-regulation. For example, there would be up to 2^50^ or ∼10^15^ possible expression states for 50 genes each with two states (on and off) if every gene expresses independently. Provided that distinct expression states can lead to similar trait alterations, the likelihood of observing similar expression states for similar trait alterations would be very low. However, the number of possible expression states would reduce dramatically if the 50 genes are subject to co-regulation (the number would be 2 for perfect co-regulation). As a consequence, the likelihood of observing similar expression states for similar trait alterations would be high. Because co-regulated gene modules must be built and/or maintained by selection, genes responsible for an unimportant trait subject to little selection are unlikely to evolve co-regulation.

**Figure 3.**
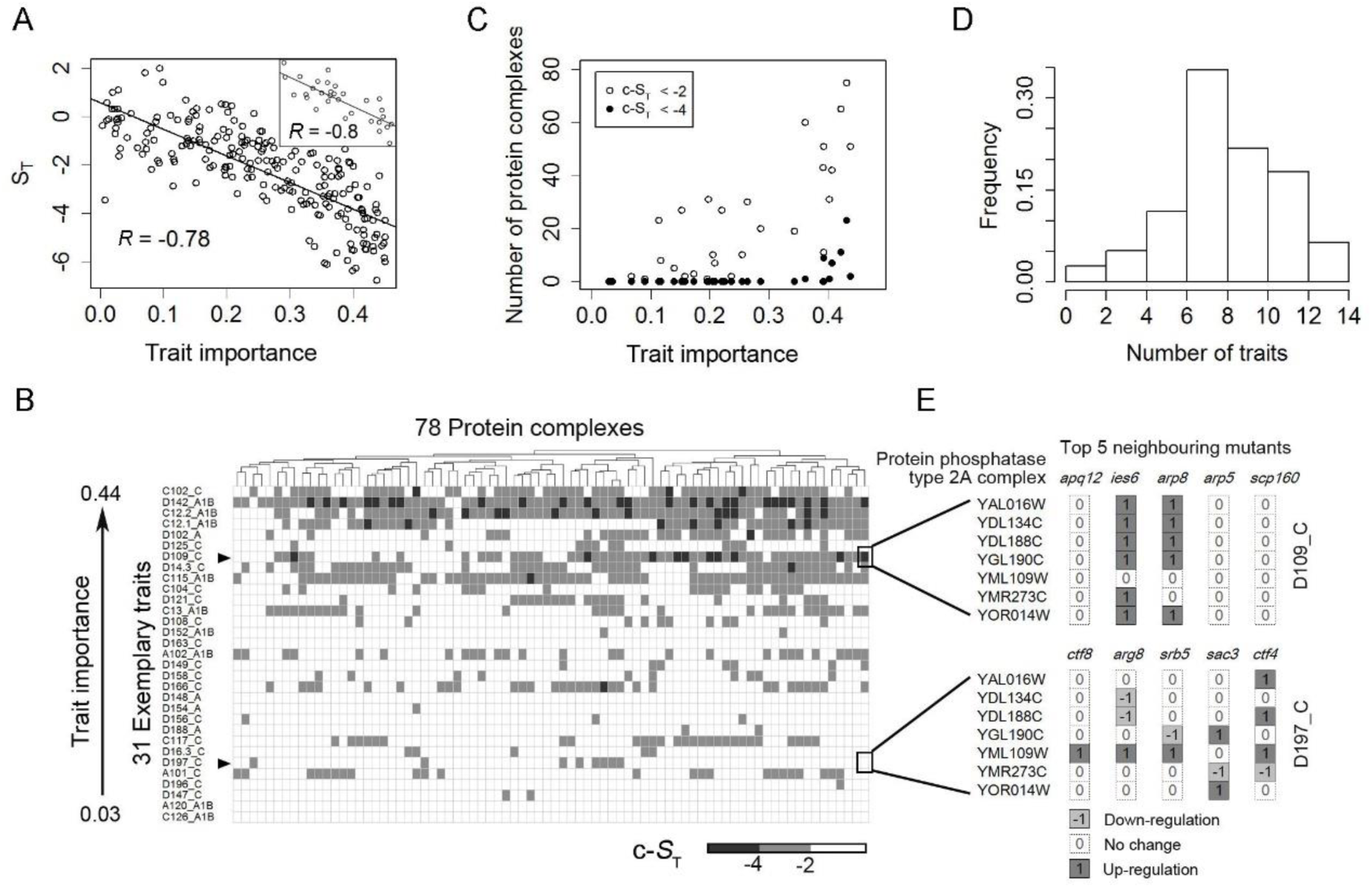
**(A)** *S*_T_ is explained by the trait’s evolutionary importance (Pearson’s *R* =-0.78, n = 216, *P* < 10^−16^). A similar pattern is shown as an inset for exemplary traits that are largely independent from each other (*R* = −0.8, n = 31, *P* < 10^−7^). **(B)** The c-*S*_T_ of 78 protein complexes in the 31 exemplary traits that are sorted by trait importance. Different traits show a different spectrum of significant c-*S*_T_ (< −2 or −4), suggesting trait-specific effects provided by the protein complexes. **(C)** More significant c-*S*_T_ are found for important traits, suggesting that the selection for maintaining the expression coordination of the protein complexes be mediated through these morphological traits. **(D)** Frequency distribution of the numbers of exemplary traits that are affected by each of the 78 protein complex (c-*S*_T_ < −2). **(E)** The expression profile of an example protein complex in the top 5 mutants of the traits D109_C and D197_C, respectively; shared expression profiles are found in neighboring mutants of D109_C (c-*S*_T_ < −4), the evolutionarily more important trait that measures the distance from neck to bud’s nucleus.

Although the simulation was conducted with arbitrary settings, it provided clues on how natural selection might be involved in shaping *S*_T_. An immediate prediction is that the yeast traits with *S*_T_ < −2 should be more important than those of −2 < *S*_T_ < 2. Following previous studies (Ho and Zhang 2014; Chen et al. 2016), we used cell growth rate as a proxy of fitness, which was reasonable for the single-celled yeast, and calculated the correlation with fitness for each of the 216 traits (Supplemental Table 2) (Materials and Methods). Traits that are strongly correlated with fitness can be regarded as evolutionarily important and subject to strong selective constraints. In support of the prediction, the *S*_T_ of a trait was well explained by the trait importance (Pearson’s *R* = −0.78, *n* = 216, *P* < 10^−16^; Fig. 3A); the pattern held when 31 exemplary traits that are less related with each other were considered (Pearson’s *R* = −0.80, *n* = 31, *P* < 10^−7^; inset of Fig. 3A) (Materials and Methods). The result cannot be explained by the varied genetic complexity of the traits (Supplemental Fig. 4), which is defined by the fraction of genes whose deletion effects are statistically significant, or the varied measuring repeatability (Supplemental Fig. 5).

**Figure 4.**
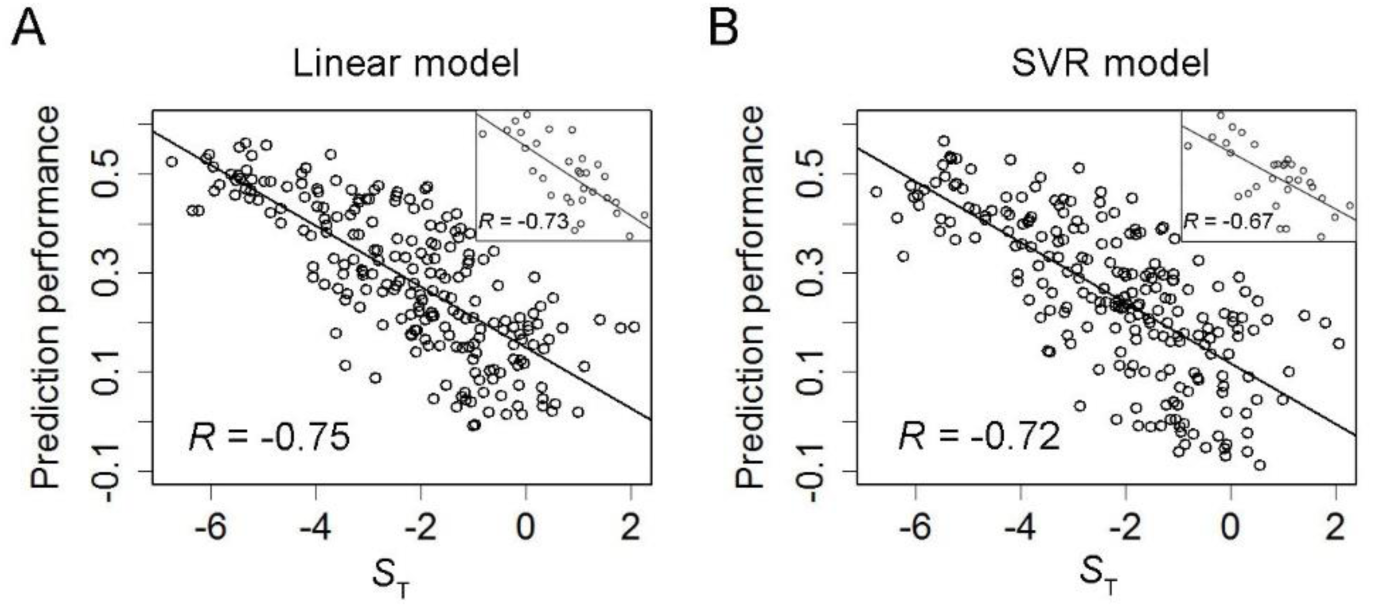
*S*_T_ underlies the prediction performance of the linear model **(A)** and the SVR model **(B)** for all the traits (Pearson’s *R* = −0.78 or −0.72, n = 216, *P* < 10^−16^) or the exemplary traits (inset) (Pearson’s *R* = −0.73 or −0.67, n = 31, *P* < 10^−5^ and *P* < 10^−4^ respectively). Here the prediction performance is measured by Pearson’s correlation coefficient between observed and predicted trait values of the testing mutants.

According to the simulation and reasoning, natural selection reduces *S*_T_ through promoting co-regulated gene modules to reduce the number of possible expression states that lead to the same phenotypic outputs. We examined the yeast genes forming protein complexes, which are known to be co-regulated (Teichmann and Babu 2002). We considered 78 core yeast protein complexes each encoded by at least five different genes (Materials and Methods) (Supplemental Table 3). Because the traits examined here represent nearly all dimensions of the yeast morphology, it is likely that, for at least some of the protein complexes, their co-regulations are due to the selective constraints on these traits. For each protein complex we computed <u>c</u>omplex-specific *S*_T_ (c-*S*_T_ for short) by using the same formula but considering only the few genes encoding the focal protein complex (Materials and Methods). Consistently, c-*S*_T_ < −2 suggested similar expression profiles of the protein complex genes given the same/similar trait alterations. If the co-regulation within a protein complex is built/maintained by selection irrelevant to the morphological traits, no correlation would be expected between c-*S*_T_ and the strength of selection on these traits. However, we found that, for many of the protein complexes, c-*S*_T_ tends to be significant (< −2 or even < −4) in important traits (Fig. 3B-C). This result suggested that natural selection acting on these morphological traits account for the co-regulations of the protein complexes genes. Notably, to achieve this, the protein complexes must causally influence the traits instead of being reactive to the traits, because selection on a trait is unable to shape the genes just reactive to the trait. There was a different spectrum of significant c-*S*_T_ even among the important traits (Fig. 3B), and also the number of exemplary traits that are presumably influenced by each protein complex varied substantially (Fig. 3D). This heterogeneity suggested that the significant c-*S*_T_ cannot be not explained by a single factor, such as the response to growth reduction. Thus, by associating known protein complexes with the many morphological traits we revealed both common and trait-specific mechanistic insights into the yeast traits (Supplemental Table 4). For example, the Isw1 complex involved in modifying chromatin structure appeared to underlie C102_C, a trait measuring the contour length of budding cell; and the TRAPP complex involved in transporting vesicles from ER to plasma membrane appeared to affect D109_C, a trait measuring the distance from neck to budding cell’s nucleus. These understandings provide rich information for future studies on cell morphology. To better explain the analysis of this part, using the protein phosphatase type 2A complex as an example we plotted the expression profiles of the 7 member genes of this complex in the top 5 mutants of an important trait D109_C and an unimportant trait D197_C, respectively (Fig. 3E).

## *S*_T_ is an indicator of recurrent patterns of an expression-trait relationship

Recurrent patterns are the prerequisite for revealing rules from empirical data (Ohnomachado 2001; Veer and Bernards 2008). Since *S*_T_ effectively measures the titer of recurrent patterns of an expression-trait relationship, it may predict how much we could possibly learn from an expression-trait relationship. This point is supported partly by the above analysis of protein complexes that were found primarily for the traits with *S*_T_ < −2. To make a more general demonstration of the issue, we may seek for analytical rather than just empirical evidence. We considered machine learning, which is often more capable than humans in dealing with complex expression-trait relationships. To ensure that the learning is mathematically tractable we studied the multiple linear regression model and gauged the learning performance using the correlation between observation and prediction. We showed analytically that *S*_T_ well approximates the learning performance (Materials and Methods, with details in Supplemental Text). To gain further empirical support we analyzed the yeast expression-trait data by machine learning. For every trait we used the same set of mutants as training data, ran the same learning processes without artificial feature selection, and tested the learned models in the same testing mutants (Materials and Methods). This design made the prediction performance for different traits comparable. Consistent with the analytical demonstration, the varied prediction performance for the different traits by linear models was well explained by *S*_T_ (Fig. 4 & Supplemental Fig. 6). We also tested a non-linear support vector regression (SVR) model. We found that the prediction performance was also highly correlated with *S*_T_ (Fig. 4 & Supplemental Fig. 6), although we were not able to prove analytically the relationship between *S*_T_ and the prediction performance for non-linear models. We noted that the predictions could be improved with more sophisticated models that can capture high-order gene interactions or with the inclusion of pre-existing knowledge. However, the strong correlations suggested that *S*_T_ be a good measure of the relative titer of recurrent patterns, which determine the success of any learning practice based on empirical data.

## Discussion

The central question addressed in this study is about the number (*N*) of different expression states that underlies a given trait abnormality. We developed a simple statistic (*S*_T_) for describing such an *N*-to-one expression-trait relationship. Using yeast data we showed that *S*_T_ is constrained by natural selection, which helps build/maintain co-regulated gene modules to reduce *N*, and that *S*_T_ well predicts the relative tractability of over 200 expression-trait relationships. Considering the similarity to the entropy of a thermodynamic system, *S*_T_ can be viewed as the expression entropy of a biological trait. Accordingly, stochastic thinking might be necessary to deal with the strong expression uncertainty of a trait with *S*_T_ ≥ 0, a proposition reminding biologists of the paradigm shift in modern physics from classical mechanics to statistical mechanics. Because the definition of *S*_T_ is independent of the yeast system and natural selection as the ultimate determinant of *S*_T_ applies to all organisms,the main conclusions of this yeast-based study are likely to be of general meaning, although further work is certainly required to test the findings in other organisms.

## Materials and Methods

### Data

The microscopic images of triple-stained yeast cells are generated and analysed by Ohya, et al. (2005), characterizing quantitatively 501 morphological traits for 4,718 different yeast mutants each lacking a non-essential gene (SCMD). A proportion of the morphological traits (220 traits) with information of individual cells are considered by Ho, et al. (2014). We consider just 216 traits that show consistent information between the two previous studies in this analysis. The 216 morphological traits are also characterized for 122 wild-type yeast cell populations by Ohya, et al. (2005). For each trait the mean and standard variation of the 122 wild-type values are computed, and the raw trait value of a mutant is then scaled into Z-score using the mean and standard deviation of the wild-types. In addition, the cell growth rate of 4,449 yeast gene-deletion mutants are measured by Qian, et al. (2012), and the microarray-based expression profile of 1,484 mutants are generated by Kemmeren, et al. (2014). Because there are 6,123 genes covered in the microarray, the cutoff of defining expression up-regulation or down-regulation relative to wild-type is set to *p* < 0.0001 (the number of expected expression changes by chance is ∼0.6). With manual corrections for the annotation of the different datasets, there are 1,345 mutants each with all the information of 216 morphological traits, growth rate, and expression profile. A total of 409 yeast core protein complexes are determined by Benschop et al. (2010), and 78 encoded by at least five different genes are considered in this study.

### Define the terms or concepts used in this study

***S*_T_ and c-*S*_T_:** The Euclidean distance of the expression profiles of two neighboring mutants (E(*M*_*i*_, *M*_*i* +1_)) is calculated as Eq. 1 for *S*_T_, where *e*_*i,j*_ is the expression state of gene *j* in mutant *i*.

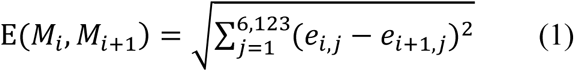

The c-*S*_T_ is computed similarly to *S*_T_ but considering only the few genes encoding the focal protein complex and using a modified Euclidean distance (E(*Mc*_*i*_, *Mci+1*)) as Eq. 2 to controls for the gene number (N*c*).

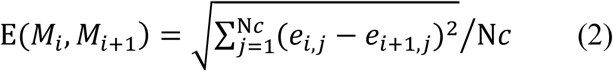

### Exemplary traits

The 216 traits are not independent, and 31 exemplary traits are derived by clustering analysis (R package, ‘apcluster’, negDistMat, *r* = 2, Bodenhofer, et al., 2011) of the 1,345 mutants with.

### Trait evolutionary importance

The Pearson’s *R* between the absolute Z-score of a morphological trait and cell growth rate is computed among the 1,345 yeast mutants. Because cell growth rate is a reasonable proxy of fitness for the single-celled yeast, morphological traits with a stronger correlation with fitness (larger *R*) are regarded as evolutionarily more important than those with a smaller *R*.

### Trait dissimilarity

To make the different traits more comparable, for each trait the average |Z_i_ – Z_i+1_| / (|Z_i_| + |Z_i+1_|) is used to represent the trait dissimilarity between neighboring mutants, where Z_i_ and Z_i+1_ are the Z-score trait value of the sorted mutant *i* and *i+1*, respectively (i = 1, 2, …, 1344).

### Genetic complexity

The genetic complexity of a trait is measured by the fraction of non-essential yeast genes that affect the trait (*f*_genes_), which is obtained from Ho, et al (2014).

### Trait measuring repeatability

Often a few hundred cells of a mutant are examined in Ohya, et al (2005). We randomly divide the individual cells of each mutant into two equal halves and compute the traits for each half separately. For each trait the Pearson’s *R* between the two halves among the 1345 mutants is considered as the measuring repeatability of the focal trait.

## Simulation of the evolution of a gene network

Suppose a network consisting of 50 worker genes regulated by four regulators and four traits each determined by a non-overlapping group of ten worker genes (remaining ten workers not associated with any trait). The value of a trait is the geometric average of the absolute expression levels of the specific ten workers underlying the trait.

To build the initial network structure we assign regulator-worker interactions by assuming equal probability for all regulator-worker pairs so that on average a regulator controls 10 workers. The regulatory strength of a regulator-worker interaction considers two parameters: a regulator-specific trans-coefficient *q*, and a cis-coefficient *r* that is specific for each regulator-worker interaction. All *q* and *r* values are randomly chosen from standard normal distribution. The relative expression of worker *j* is then given by Eq. 3, where c=2is a constant baseline expression level for every worker, R is the number of regulators that control worker *j*.

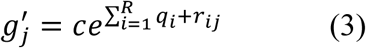

Assuming a simple resource allocation model for expressions, the absolute expression level of worker *j* is given by Eq. 4, where *C* = 100 is the total expression capacity (only workers’ expressions are considered).

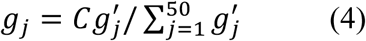

With this network structure unaltered, we simulate 100 scenarios each with different *q* and *r* values to derive the initial trait values. The average of the 100 values is used as the expected initial value *t’* of a trait. We define the optimal value of trait *k* in environment *l* by Eq. 5, where *d* acts as the “direction” factor to determine the direction of selection for optimality, and *s* is a random number of the uniform distribution between –1 and 1, which provides additional variability for the optimal trait values in different environments.

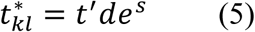

A total of ten environments are considered, and, for each trait, *d* equals to 100 in five environments and to 0.01 in the other five environments. In addition, for the ten environments considered, the trans-coefficient *q* of a regulator may vary across different environments (for each regulator there are different *q* values drawn from standard normal distribution in the initial network), but the cis-coefficient *r* of a regulator-worker interaction remains constant across the environments.

The fitness of a specific network in environment *l* is calculated as Eq. 6, where *w*_*k*_ is the fitness weight for trait *k* (*w*_*1*_ = *w*_*2*_ = 0.1, *w*_*3*_ = *w*_*4*_ = 10), and *t*_*kl*_ is the value of trait *k* in environment *l*.

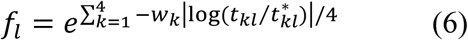

The combined fitness of a network is the harmonic average of the ten environment-specific fitness.

To simulate the evolution of the network we consider a population of 100 individuals. For each individual, four kinds of mutations are applied independently each with 10% probability: (i) adding a random number drawn from the uniform distribution between –0.1 and 0.1 to a random *q*_*il*_; (ii) adding a random number drawn from the uniform distribution between –0.1 and 0.1 to a random *r*_*ij*_; (iii) creating an interaction for a random regulator-worker pair; and (iv) deleting an random regulator-worker interaction. After the mutations are applied, the new fitness of each individual/genotype is calculated. The 100 individuals/genotypes of the next generation are obtained by drawing with replacements randomly from the current 100 individuals/genotypes according to the new relative fitness of the individuals/genotypes. This process contains mutation, selection and drift, and is repeated for 10,000 generations.

## Analytical demonstration of S_T_ as an approximation of the machine learning performance (with details in Supplemental Text)

For a multiple linear regression model, the response vector 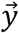 is formulated by feature matrix *X* as (7):

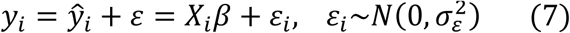
where *y*_*i*_ and 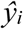 are the observed and predicted value of a response variable in *i*th sample, and *X*_*i*_ denotes the *i*th row of *X*. According to the routine training-testing paradigm with *L*-2 regularization in machine learning, *β* is analytically estimated by (8), and *λ*_*opt*_ is an optimal hyper-parameter determined by cross validation.

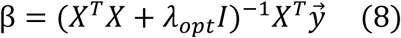

*R*^2^, which is square of Pearson’s *R* in a linear regression analysis, is given by (9) when data is normalized before learning.

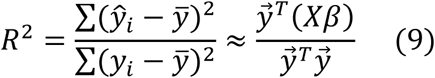

The denominator 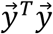 in (9) can be assumed to be constant under normalization when the sample size is fixed. Thus, *R*^2^ is determined (10).

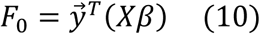

Combining (8) with (10), we get:

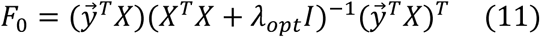

By singular value decomposition (SVD), 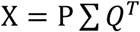, where *P* and *Q* are orthogonal matrices. Then *F*_*0*_ can be rewritten as:

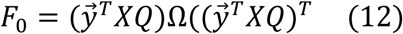
where Ω = diag 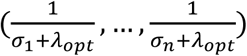, and *σ*_*i* (*i* = 1, 2, …, *n*)_ is the square of ith singular value of X. With a common *λ* larger than the maximum of *σ*_*i*._ The final form of *F*_*0*_ can be approximated by (13) based on Talyor expansion (details in Supplement Text).

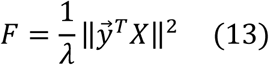

*F* can be reversely determined further to *F*^*r*^ based on rearrangement inequality as (14) (details in Supplemental Text):

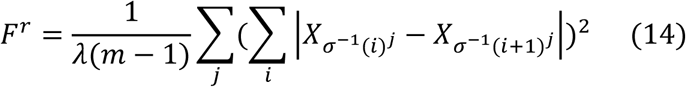
where *m* is the sample size and j =σ^-1^*(i)* represents the *j*th term of vector *y* ranked *i*th when sorted ascendingly. Finally, we can get the approximate relationship between *F*^*r*^ and *S*_T_ as (15) since the arithmetic mean is less than and approximate to the square average (details in Supplemental Text):

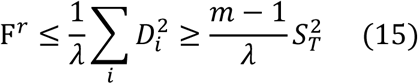
where *D*_*i*_ is the Euclidean distance between 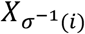 and 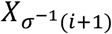.

## Empirical performance of machine learning

Supervised machine learning is conducted to infer an expression-trait function for each of the 216 traits using the 1345 mutants. The mutants are divided into a training set and a testing set with a ratio of 4:1. For each trait, the input features are all genes’ expression levels and the output is the corresponding trait value. A linear model and the non-linear support vector regression (SVR) model are considered. A ten-fold cross validation for the linear model or a five-fold cross validation for the SVR model is first carried out in the training set to tune the hyper-parameters; normal learning processes are then run in the same training set. The trained models are tested in the testing set. For each trait the performance of the trained models is evaluated by Pearson’s *R* between observed and predicted trait values, normalized root mean squared error (NRMSE), and mean absolute error (MAE), respectively. Below are the details of the two models.

### Multiple linear model

The hypothesis of multiple linear model is represented as (16), where X is an m by n matrix of expression levels, y is an m by 1 vector of trait values (m is the number of samples considered and n is the number of features), β is the parameters to be learned from training set and is defined as the minimizer of the cost function of (16), as shown by (17), where λ is the controller of contribution on the cost function by the regularization term.

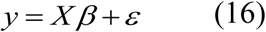

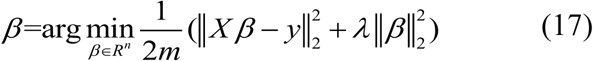

Gradient descent algorithm written in conventional MATLAB code is used to estimate β under the optimal λ determined by cross validation. Then performance of the learned model with estimated β is tested in the testing set.

**Support vector regression (SVR) model:** Epsilon-support vector regression is a non-linear regression model through kernel transformation of original features (Pedregosa et al., 2011). The hypothesis of SVR is represented as (18) with Gaussian (RBF) kernel shown as (19), where N, X and *x*_*i*_ represents the number of support vectors, original m by n feature matrix and a support vector, respectively.

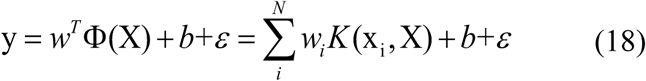

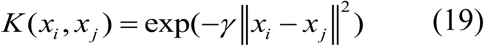

SVG solves a primal optimization problem given as (20).

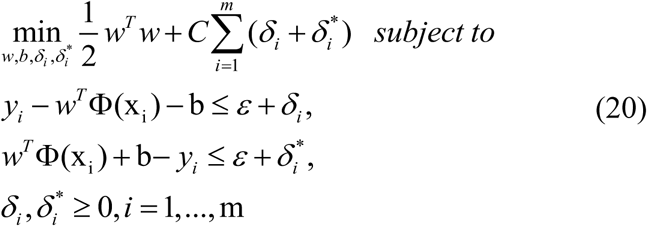

The SVR function with sequential minimal optimization (SMO) algorithm embedded in the cikit-learn package of Python achieves optimization process above. Two hyper-parameters, C and γ is tuned by grid search approach combined with a five-fold cross validation. Learned model in training set is evaluated in testing set.

## Supplemental Information

**Supplemental text** - The theoretical analysis of the connection of *S*_T_ and *R*^2^

The mathematical essence of predicting phenotype based on gene expression profile with linear model is conducting a multiple linear regression from feature matrix *X*_m×n_ to response vector 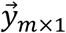 The model is commonly expressed as (1).

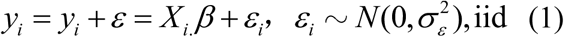
where *y*_*i*_ and 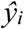 are the observed and predicted value of response variable in *i*th sample, and *X*_*i*_ denote the *i*th row of *X*. According to the routine training-testing paradigm with *L*-2 regularization in machine learning, *β* is analytically estimated by (2).

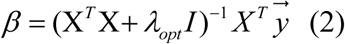
where *λ*_*opt*_ is the optimal hyper-parameter, determined by cross validation.

To evaluate the performance of a model, *R*^2^ called coefficient of determination in regression analysis, or the square of multiple correlation coefficient (Pearson’s *R* between observed and predicted value), is often used, given by (3).

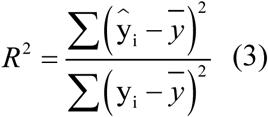

For convenience, feature variables and response variable are usually normalized before learning since the normalization process does not change prediction results. Thus, *R*^2^ can be approximated by (4) due to normalization and the orthogonality of 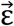 and 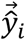.

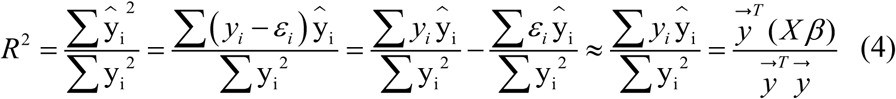

The denominator 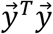 (4) can be approximated to be constant under normalization and fixed sample size. Thus, *R*^2^ is approximated by *F*_0_ given by (5).

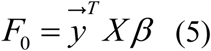

Combining with (2), we get (6):

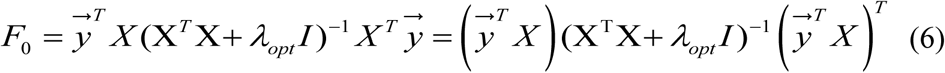

By singular value decomposition (SVD), we get

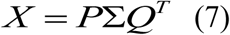
where ∑ is *m* by *n* matrix with singular value in the diagonal positions, and *P* and *Q* are orthogonal matrices satisfying

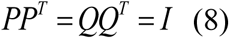

Thus,

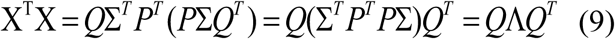
where Λ diag(σ_1_…σ_n_) satisfying σ_1_≥…≥ σ_n_ ≥0 in which σi is the square of jth largest singular value.

Thus,

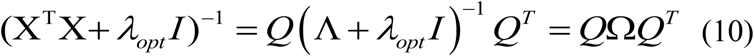
where

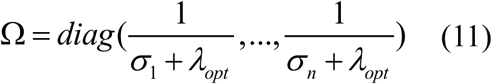

A common *λ*, which is larger than *σ*_1_, is used in place of *λ*_*opt*_. Based on Taylor expansion,

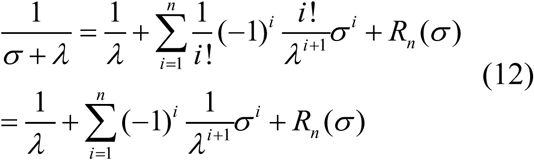

Combine (6) - (12), we get (13) (data not shown),

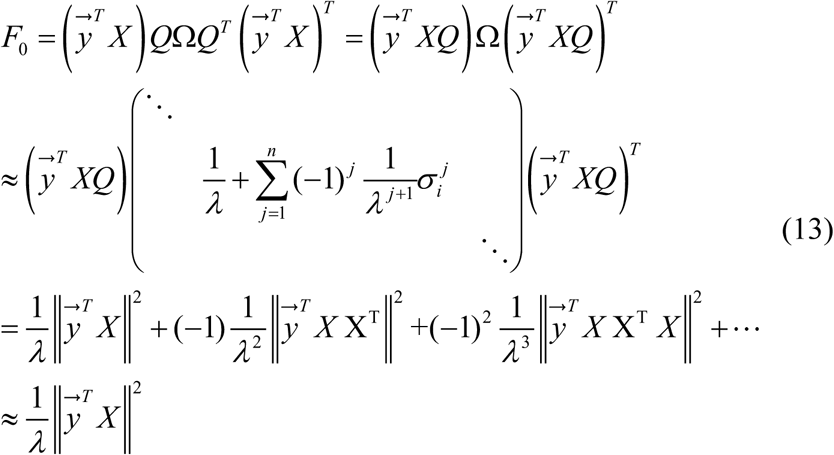

Thus, *F*_0_ is approximately determined by a quadratic form given by (14), denoted as *F*.

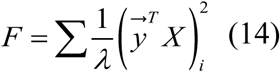

Equation (14) gives us another elegant predictor independent of learning process. However, its meaning is hard to be interpreted in empirical science.

Then, set

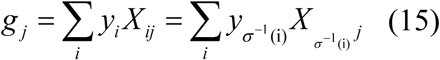
wherej *= σ*^-1^(i)represents that the *j*th term of vector *y* ranks *i*th when sorted from smallest to largest.

And we can get

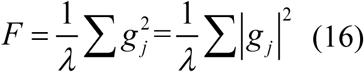

We can further deduce from (16) according to the logic below. Set *λ* fixed and *F* is maximum when each of absolute *g*_j_ is maximum. In terms of *y*, absolute *g*_j_ is maximum when *X*_.j_ has the same or reverse order with *y* according to Rearrangement Inequation, which is further approximately reversely equivalent to a form given by (18).

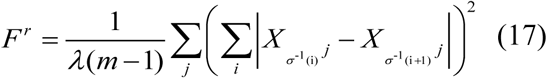
where *σ*^-1^(*i*), defined by a response variable, is consistent with the previous description.

Based on the fact that the arithmetic mean is less than the square average, we can find that (17) has a form consistent with arithmetic mean, and our *S*_T_ defined based on biological logic has a form consistent with arithmetic mean, too. Here, we build the approximate relationship between *F*^*r*^ and *S*_T_ by seeking a common quadratic form as their common upper limit, given by (18).

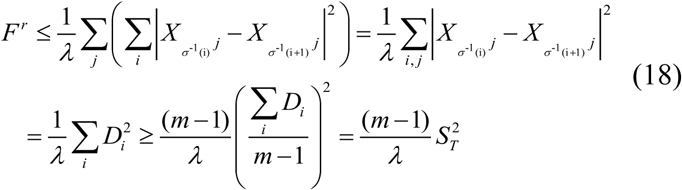
where *m* is the sample size and *D*_*i*_ is the Euclidean distance between 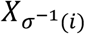 and 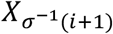. Then we set

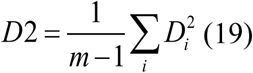

*D*2 defined in (19) is just the common quadratic form and is also a predictor. *S*_T_, which is a different form of *F* or *F*^*r*^, can be better interpreted as the titer of patterns underlying a response variable. In our circumstance, smaller *S*_T_ means smaller expression distance between neighboring trait values, which means less recurrent patterns underlying a focal trait, and better tractability of the trait.

## Some remarks

*1. β* in the proving is an approximation of *β*_train_ which is determined in training set.

2. In practice, the performance is evaluated in testing set due to the overfitting in train set, because *λ*_*opt*_ is also an approximation by cross validation. In our deduction, we assumed *λ*_*opt*_ is the truly optimal parameter and *R*^2^ can be evaluated in the total set.

3. In the deduction, we have used a common *λ* which is larger than the square of the largest singular value in place of the *λ*_*opt*_. The approximation between the prediction based on *λ* and that based on *λ*_*opt*_ is shown in the following figure.

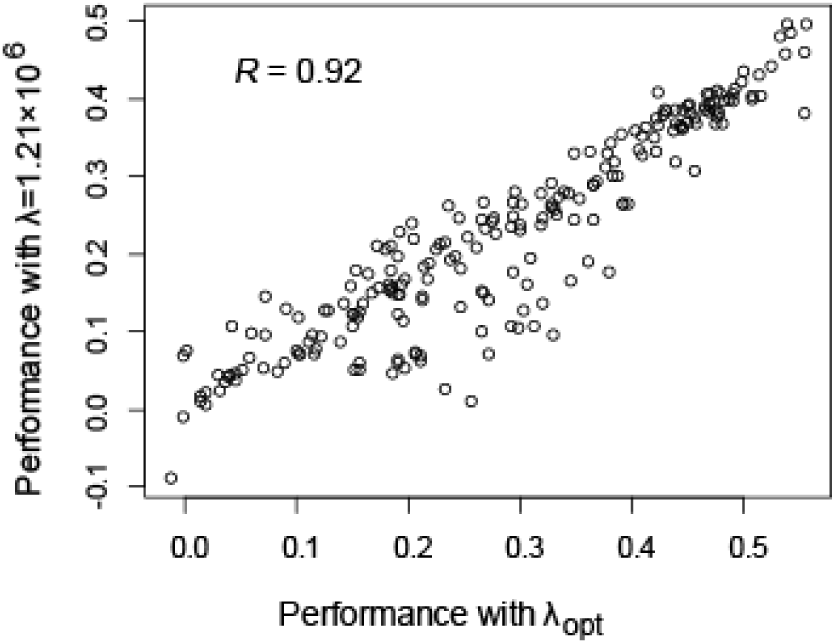

In the above figure, the *x*-axis and the *y*-axis are Pearson’s *R* between observed and predicted trait values with λ_*opt*_ and common λ equal to 1.21×10^6^, respectively. The maximum of *σ*_*i*_ is less than 1.21×10^6^. The approximation between two results are supported by the high correlation between them (Pearson’s *R*=0.92, *P* < 10^−16^).

4. Reverse approximation between *F* and *F*^*r*^

A sequence is an ordered set {*a*_i_} and the sum of neighboring distances is calculated by 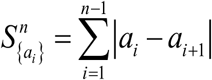 for an *n*-element ordered sequence. The minimal value of 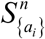 is denoted as 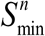 which is obtained when {*a*_i_} is organized in a monotonous order. For example, we can get 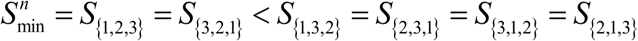. A sequence ordered from smallest to largest or from largest to smallest is called complete ordered sequence and denoted as 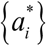. For convenience, 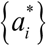 is referred to be ordered from smallest to largest below by default since the reverse ordering is equivalent. Here we want to prove the equivalence relationship between 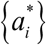 and 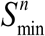. In other words, we want to prove 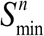 is obtained if and only if {*a*_*i*_} is ordered as 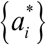.

We first map {*a*_i_} onto a number axis, in which each point corresponds to a number in {*a*_i_} and arranged from the smallest to the largest. 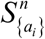 is equivalent to the track length defined as the sum of corresponding segment length on the axis of a track, plotted outside the axis and through every number according to the order in {*a*_i_} without circles. In the track, each number has a degree of 2 except the first and the last which have a degree of 1, and all the numbers are on the same track, which are two properties of the track not constrained. Thus, 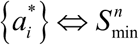 is equivalent to that the track length is minimal in the situation of continuously joining from the smallest to the largest.

(i). 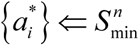 with mathematical induction:

When {*a*_i_} has two elements and a_1_*≤* a_2_ 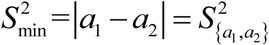. So 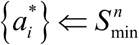. Assuming that 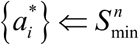 for *n ≤ k*, we can get 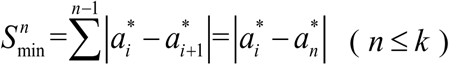. [(n *≤* k). So we can derive that continuously joining according to 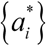 gives the minimal track length for n *≤* k. Next, we will prove 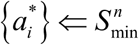 is satisfied for *n=k +1* with proof by contradiction. Assuming that 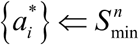 is not satisfied for *n=k +1*, therefore, there must exist a track whose track length is less than the track from the smallest to the largest defined in 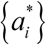. There must exist repeated covering without any vacancy intervals which are defined as intervals not covered by any corresponding segment of the track. Otherwise, the track is just the track defined in 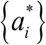 which has no repeated covering and vacancy intervals,too, which produces contradiction. When the track has repeated covering, it can’t has a track length less than the track defined in 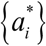 and thus produce contradiction. Thus, there must exist at least a vacancy interval. Based on assumption for n *≤* k, the two parts separated by this interval must be ordered as 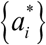. Thus, there must exist four number, the first and the last and the two on both sides of this interval. This is contradicted to the track property that all the numbers are on the same track. Therefore,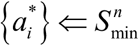 is satisfied for *n = k* +1and further any *n*.

(ii) 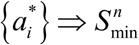

All tracks must cover the whole segment from the smallest to the largest, which is obvious. Thus, the length of any track must be no less than track length of 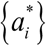 which only covers one time for each sub-segment of the whole segment. Then, 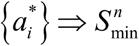.

## Legends of supplementary tables

**Supplemental Table 1** Identity of the 1,345 gene-deletion mutants.

**Supplemental Table 2** Characterized features of the 216 morphological traits, including *S*_T_, trait dissimilarity, evolutionary importance, measuring repeatability, genetic complexity, prediction performance by linear model and by SVR model, respectively, being exemplar or not, and the general trait description.

**Supplemental Table 3** Information of the 78 protein complexes.

**Supplemental Table 4** The c-*S*_T_ of the 78 protein complexes in each of the 216 traits.

## Supplemental Figures

**Supplemental Fig. 1.**
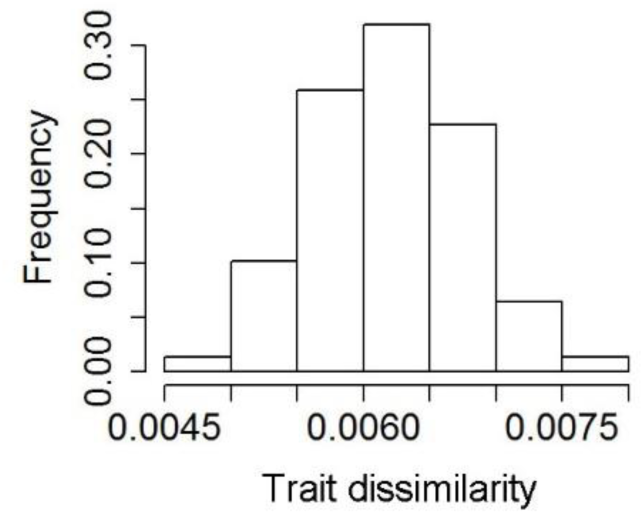
Highly similar trait values between neighbouring mutants. The average trait dissimilarity between neighbouring mutants is shown at the x-axis for the 216 yeast traits.

**Supplemental Fig. 2.**
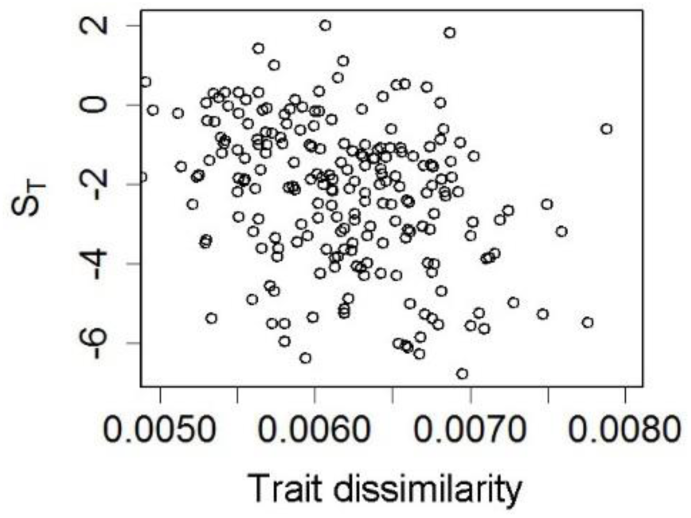
Traits with large *S*_T_ (> −2) do not show strong trait dissimilarity between neighbouring mutants. The correlation between *S*_T_ and trait dissimilarity is even slightly negative (Pearson’s *R* = −0.32, n = 216, *P* < 10^−5^).

**Supplemental Fig. 3.**
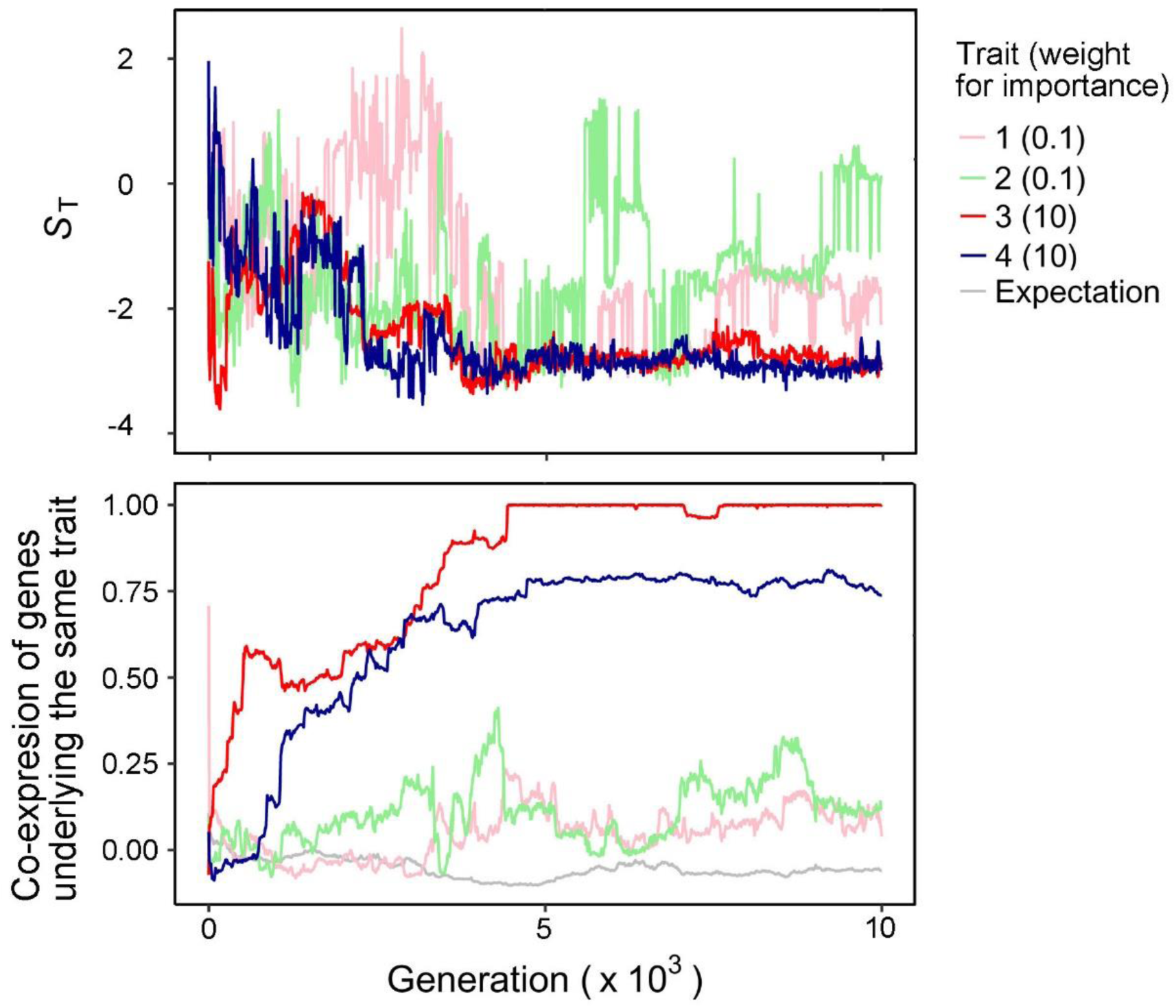
With the course of evolution the reduction of S_T_ and the increase of co-expression are stronger for important traits (red and blue lines) than unimportant traits (green and pink lines). The co-regulation/co-expression of genes underlying the same trait is measured by the average Pearson’s *R* of the expression levels in the ten environments among the genes. The gray line shows the Pearson’s *R* calculated using all gene pairs that do not affect the same trait. The S_T_ and *R* are calculated and plotted every 10 generations.

**Supplemental Fig. 4.**
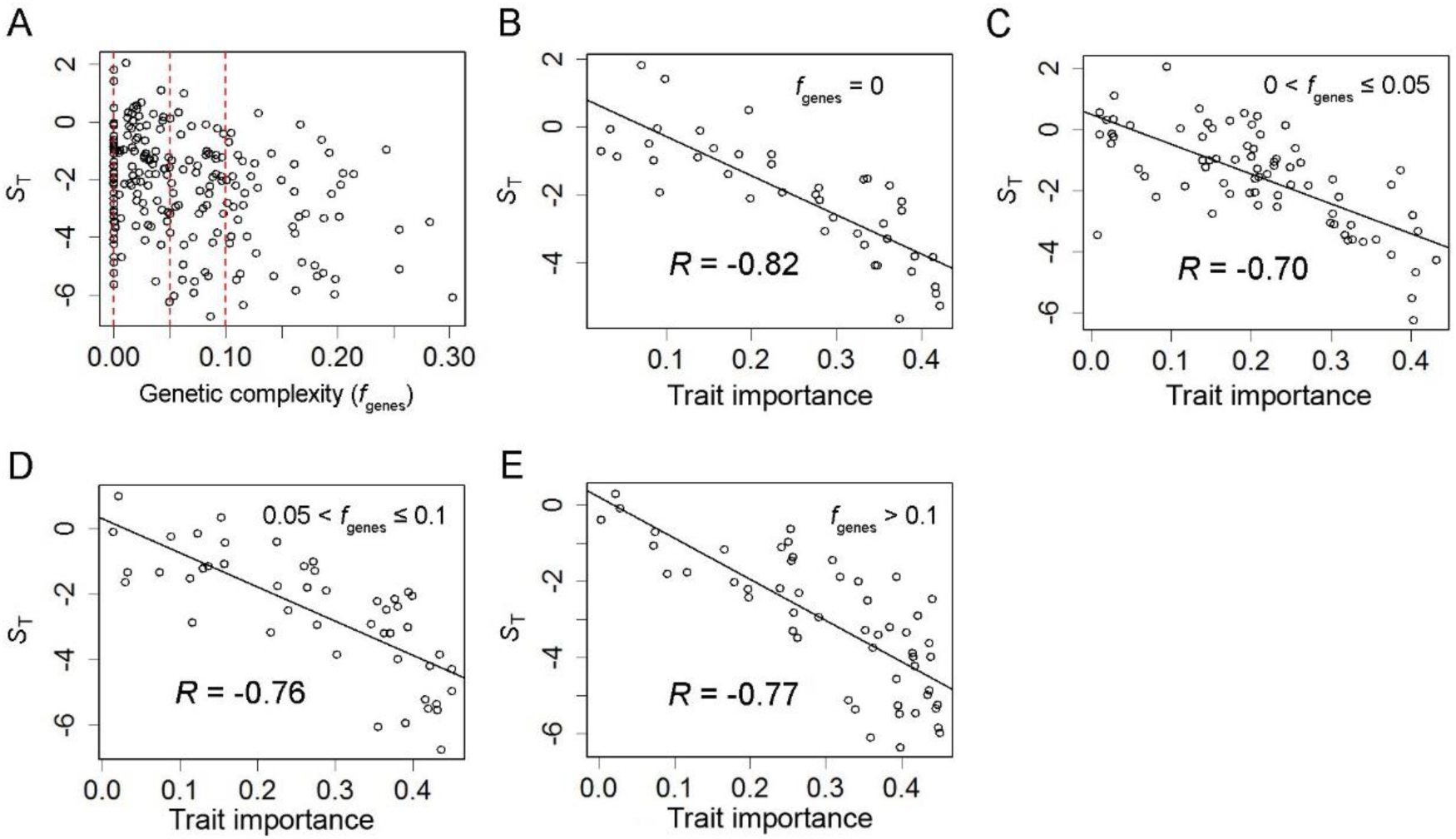
Genetic complexity measured by *f*_genes_ does not account for the relationship between *S*_T_ and trait importance. **(A)** There is a slightly negative correlation between *S*_T_ and *f*_genes_ (Pearson’s *R* = −0.31 n = 216, *P* < 10^−5^). The three red dashed lines mark *f*_genes_ = 0, 0.05, and 0.1, respectively. The strong negative correlation between *S*_T_ and trait importance holds for traits with different levels of genetic complexity: **(B)** *f*_genes_ = 0; **(C)** 0 < *f*_genes_ ≤ 0.05; **(D)** 0.05 < *f*_genes_ ≤0.1; **(E)** *f*_genes_ > 0.1.

**Supplemental Fig. 5.**
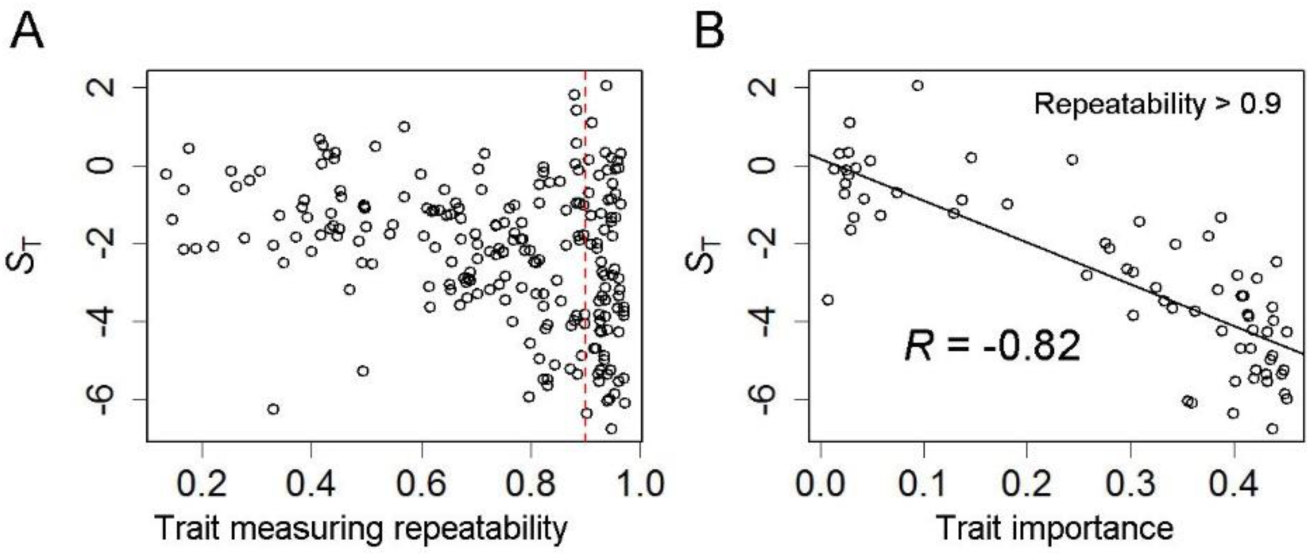
Trait measuring repeatability does not account for the relationship between *S*_T_ and trait importance. **(A)** There is a weak negative correlation between *S*_T_ and trait measuring repeatability (Pearson’s *R* = −0.31, n = 216, *P* < 10^−5^). The red dashed line marks the measuring repeatability of 0.9. **(B)** The strong negative correlation between *S*_T_ and trait importance hold for the 67 traits with good measuring repeatability (> 0.9).

**Supplemental Fig. 6.**
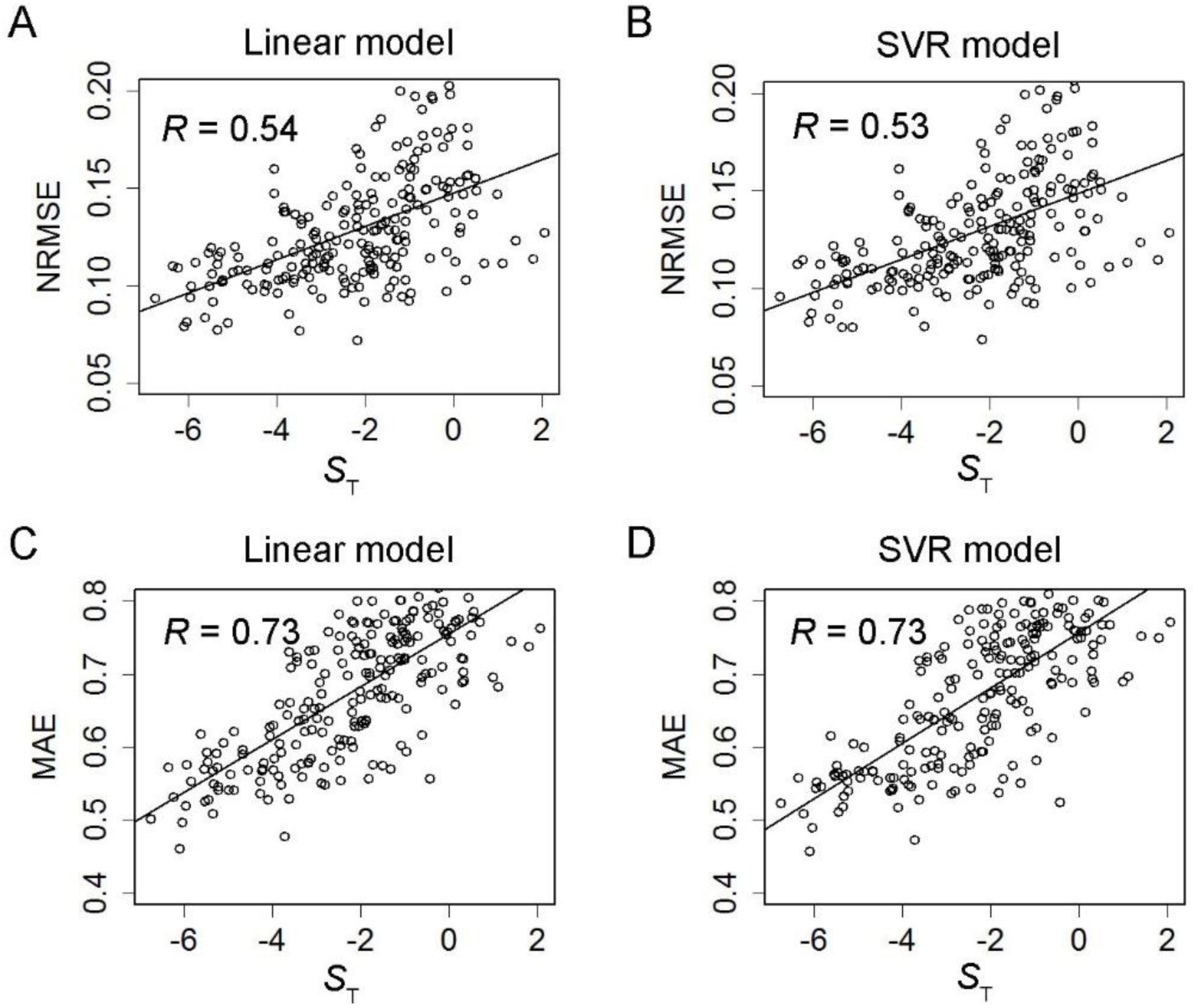
Same as Fig. 4, except that the prediction performance is measured by the normalized root mean squared error (NRMSE) or the mean absolute error (MAE) between observed and predicted trait values of the testing mutants (Pearson’s *R* = 0.54, 0.53, 0.73, and 0.73 respectively, n = 216, *P* < 10^−16^ in all cases).

